# Hippocampal-guided reconstruction of an event’s prior temporal context

**DOI:** 10.1101/2025.08.05.668710

**Authors:** Futing Zou, J. Benjamin Hutchinson, Brice A. Kuhl

**Author notes:** Co-senior author (J. Benjamin Hutchinson, Brice A. Kuhl).

## Abstract

Leading theories of episodic memory argue that when events from the past are remembered, temporally-adjacent events are also reinstated (‘temporal context reinstatement’). However, direct evidence for temporal context reinstatement is surprisingly limited. Here, we tested for temporal context reinstatement in a human fMRI continuous recognition memory experiment in which natural scene images were repeatedly encountered. For the original encounter with each scene, we defined its temporal context as the visual content of temporally-adjacent scenes. Using voxelwise encoding models, we tested whether fMRI activity patterns evoked when a scene was re-encountered carried information about the original encounter’s temporal context. Indeed, we found robust temporal context reinstatement within high-level visual cortex (lateral occipitotemporal cortex; LOTC), despite the fact that reinstated content was entirely incidental to task demands. However, temporal context reinstatement only occurred when stimuli were successfully recognized, indicating that reinstatement was behaviorally-relevant, even if incidental. Finally, the strength of temporal context reinstatement in LOTC was predicted by the similarity of hippocampal activity patterns across encounters, demonstrating distinct, but complimentary, roles for the hippocampus and neocortex in reinstating temporal context information.

## Introduction

Memories are not encoded in isolation. Rather, they are encoded in the context of other recent experience. This idea is formalized in influential theories of episodic memory which argue that events from the immediate past linger in mind as new events are encoded^1–3^. These lingering traces of past events comprise the “temporal context” into which new events are encoded. An important implication of temporal context theories is that when an individual event is later remembered, the broader temporal context of that event will be reinstated. In other words, remembering one event involves reinstatement of other events that were temporally-adjacent to that event. To date, however, direct evidence for temporal context reinstatement is surprisingly limited.

Initial inspiration for temporal context theories came from behavioral studies of free recall^4–6^. In free recall paradigms, subjects study lists of stimuli and then freely recall those words in any order they choose. Despite the lack of constraint on recall order, there is a well-established tendency for subjects to cluster stimuli during recall according to the order in which stimuli were encoded (a ‘temporal contiguity effect’)^1,3^. This phenomenon strongly suggests that recalling a given stimulus reinstates the temporal context of that stimulus—including temporally-adjacent stimuli—thereby explaining why temporally-adjacent stimuli tend to be clustered in free recall. Importantly, however, temporal contiguity effects only provide indirect evidence of temporal context reinstatement. That is, reinstatement is inferred from this behavioral phenomenon, but not directly observed.

Another line of evidence which provides more direct evidence for temporal context reinstatement comes from intracranial electrophysiological studies which show that when a stimulus is remembered, patterns of neural activity ‘match’ activity patterns that were temporally-adjacent to the initial encoding of that stimulus^7–10^. While this provides very compelling evidence that *something* is being reinstated, it is unclear what, exactly, is being reinstated. For example, temporal context theories argue that temporally-adjacent *stimuli* become part of an event’s temporal context and that these temporally-adjacent stimuli are reinstated. While reinstatement of temporally-adjacent neural activity patterns *may* reflect reinstatement of temporally-adjacent stimuli, it is impossible to resolve this issue without an explicit model of the information (content) that is carried by the reinstated neural activity patterns. Understanding whether temporally-adjacent stimuli are reinstated is important because it has potential implications for the behavioral consequences of reinstatement— for example, memory strengthening, weakening or integration^11–16^.

To the extent that temporal context reinstatement occurs, it is potentially supported by multiple, coordinated brain regions. While the hippocampus is widely thought to play a central role in temporal context reinstatement^17–19^, it may do so by mediating reinstatement within content-sensitive neocortical areas. Indeed, in other domains of episodic memory, there is compelling evidence for a division of labor between hippocampal pattern completion mechanisms and neocortical reinstatement effects^12,20,21^. Thus, establishing more direct evidence for temporal context reinstatement also requires greater specificity about the relative contributions of hippocampal versus neocortical regions.

Here, we sought to establish direct evidence for temporal context reinstatement in the human brain using data from a massive human 7T fMRI dataset (N = 8, 30-40 fMRI sessions per subject). The experimental task performed by subjects was a simple continuous recognition task in which natural scene images were repeatedly encountered (first encounter = E1; second encounter = E2). We predicted that when a stimulus was re-encountered (E2), high-level visual cortical areas would reinstate the temporal context of the *original encounter* with that stimulus. Specifically, we predicted that activity patterns evoked by E2 would carry information about the content of scene images that were *temporally-adjacent* to E1. Critically, we also predicted that temporal context reinstatement in visual cortical areas would be guided by the hippocampus^17–19^.

## Results

During fMRI scanning, subjects performed a continuous recognition memory task which required, on each trial, indicating whether a natural scene image was “old” or “new.” Behavioral memory performance for this dataset has previously been described^22^. Here, we focused on the first two encounters with each stimulus (E1, E2). Importantly, we operationalized the temporal context of E1 as the content of temporally-adjacent visual scenes (E1-1 and E1+1; “temporal context images”).

To measure temporal context reinstatement, we implemented an inverted encoding model approach in which fMRI activity patterns elicited by E2 were used to reconstruct the visual content of the E1 temporal context images (**Fig. 1**). Specifically, we extracted visual features of images from the Contrastive Language-Image Pre-Training (CLIP) image encoder (ViT-32 transformer backbone^23^)—a deep neural network that captures high-level visual representations in the human brain^24^. We then performed principal component analysis (PCA) on the extracted CLIP image embeddings and used the top 20 principal components (PCs) to define the information contained within each image. Next, we constructed voxel-wise encoding models to learn a direct linear mapping from the PCs (image content) to fMRI activity patterns. The encoding models used cross-validated ridge regression, separately for each subject. For each held-out fold, we inverted the model to reconstruct the image PCs from fMRI activity patterns. Reconstructed PCs were compared to actual PCs using cosine similarity.

**Fig 1.**
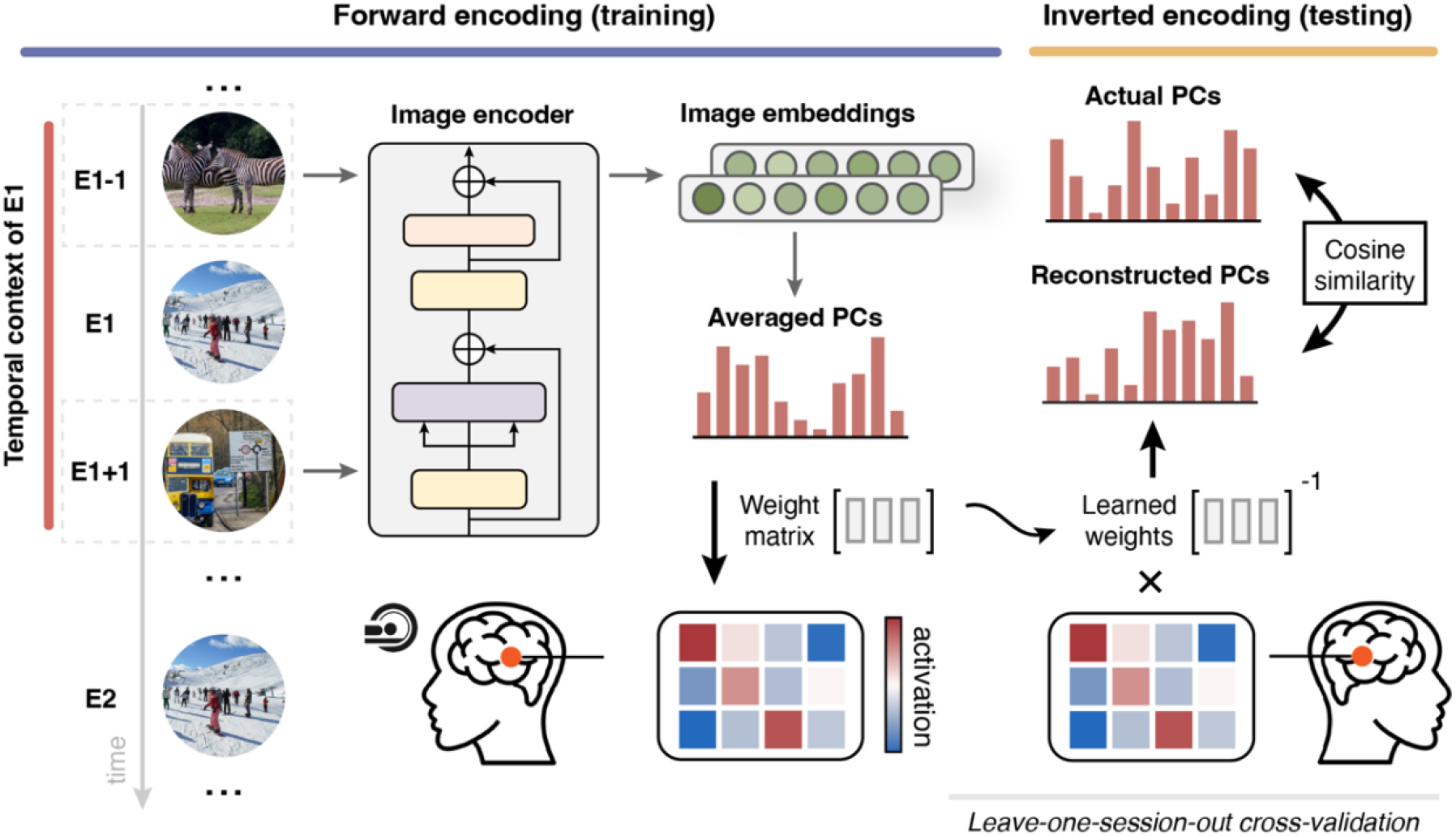
An fMRI inverted encoding model for temporal context reconstruction. We operationalized the temporal context of a stimulus’ original encounter (E1) as the content of temporally-adjacent visual scenes (E1-1 and E1+1). Scene content was measured by extracting embeddings from the CLIP visual encoder. PCA was then applied to identify the top 20 PCs. PC scores were averaged across the E1-1 and E1+1 images and these average scores (E1’s temporal context) were used in voxel-wise encoding models to learn a linear mapping to fMRI activity evoked at E2. The encoding models used ridge regression with leave-one-session-out cross-validation, separately for each subject. For each held-out fold, the encoding model was inverted such that the image PC scores were reconstructed from the fMRI activity patterns. Reconstruction accuracy was evaluated by computing the cosine similarity between the reconstructed and actual image PC scores.

We tested for temporal context reinstatement in three *a priori* regions of interest (ROIs) that are known to support memory reinstatement across different representational levels (**Fig. 2b**): angular gyrus (AG)^25^, lateral occipitotemporal cortex (LOTC)^26^, and early visual cortex (V1)^27^. As a control region, we also included primary motor cortex (M1). Our initial analyses were restricted to E1/E2 stimuli that were (1) correctly identified as “new” at E1 (correct rejection) and correctly recognized as “old” at E2 (hit) and (2) were encountered in the same fMRI session, but in different scan runs.

**Fig 2.**
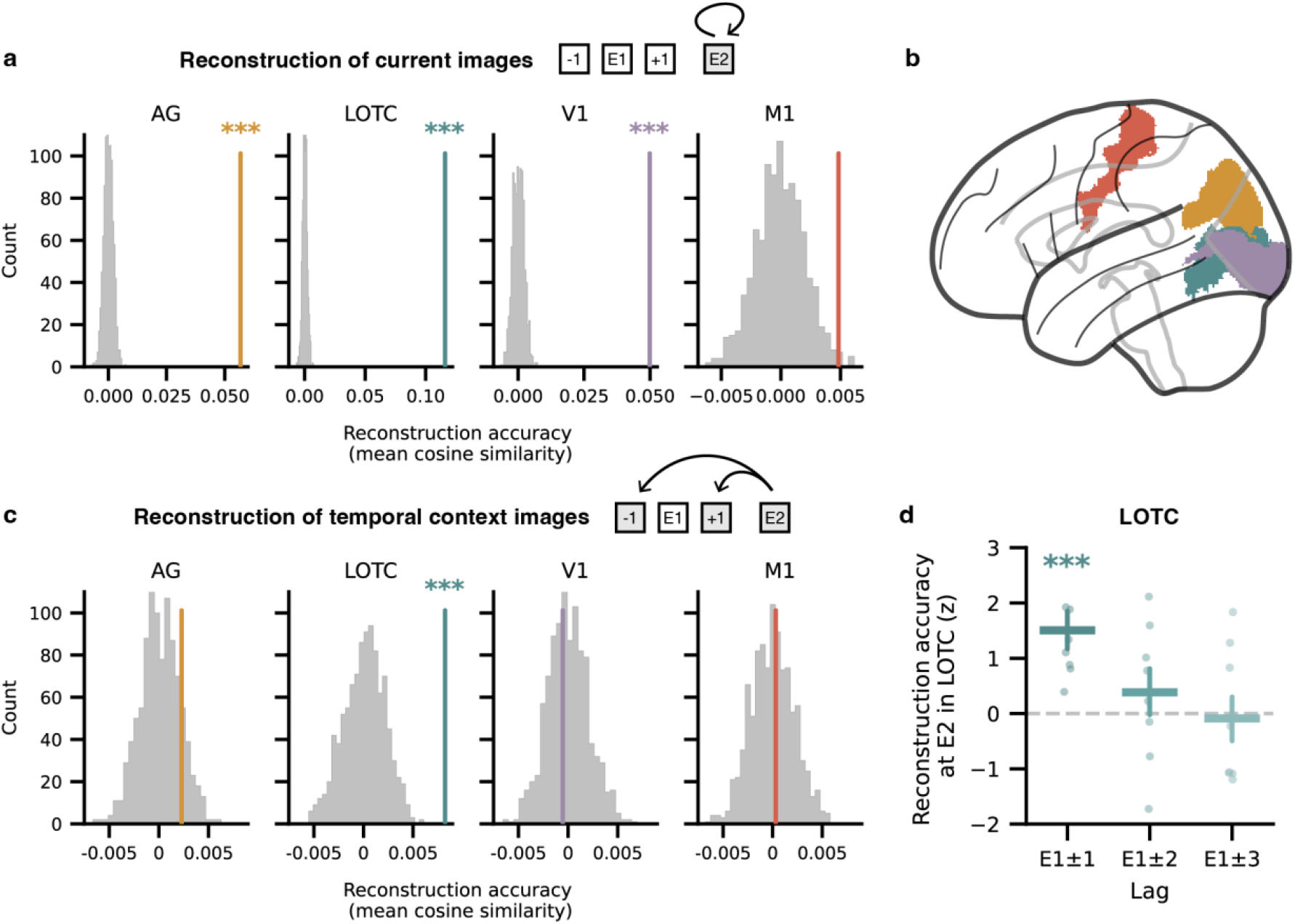
Temporal context reconstruction in LOTC. **a**, Reconstruction accuracy (from inverted encoding models) for PC scores of current images (E2) from fMRI activity patterns at E2. Reconstruction of current images was above chance in AG, LOTC, and V1 (*P*s < 0.001), but not in M1 (*P* = 0.07). Statistical significance was determined by permutation test (shuffling reconstructed image PC scores within subjects to generate a null distribution of the group-level average similarity between actual and reconstructed scores). **b**, *A priori* ROIs included bilateral angular gyrus (AG; yellow), lateral occipitotemporal cortex (LOTC; green), early visual cortex (V1; purple), and primary motor cortex (M1; red). **c**, Reconstruction accuracy of temporal context images (E1-1 and E1+1) from fMRI activity patterns at E2. Reconstruction of temporal context images was above chance in LOTC (*P* < 0.001), but not in AG, V1, or M1 (*P*s > 0.15). In **a** and **c**, density plots show the across-subject mean of the observed reconstruction accuracy (colored vertical line) relative to the null distribution (gray histograms; 1,000 iterations). Reported P-values are Bonferroni corrected to account for multiple comparisons. **d**, Temporal context reconstruction accuracy in LOTC decreased as a function of the temporal context images’ distance to E1 (*F*_2,14_ = 3.79, *P* = 0.048; repeated-measures ANOVA). Dots represent effects for each of the 8 subjects (*z*-scores from subject-specific permutation tests; see Methods) and error bars depict s.e.m.; \*\*\**P* < 0.001.

Before testing for temporal context reinstatement, we first validated our approach by testing whether fMRI activity patterns at E2 contained information about the current image (E2). In line with prior work^24^, we found that visual content information was robustly reflected in AG, LOTC, and V1 (**Fig. 2a**; permutation tests: *P*s < 0.001, corrected), but not in M1 (*P* = 0.07, corrected). These results confirm that the ROIs were sensitive to content within natural scene images and validate our approach for reconstructing image content from fMRI activity patterns.

Of critical interest was whether the fMRI activity patterns at E2 contained information about images that were temporally-adjacent to E1 (PCs from the E1 temporal context images). Importantly, because we only analyzed fMRI data from E2, we avoided any issues related to fMRI autocorrelation across E1 and E1 temporal context images. We found that information within the E1 temporal context images was successfully reconstructed from E2 activity patterns in LOTC (**Fig. 2c**; permutation test: *P* < 0.001, corrected), but not in AG or V1 (*P*s > 0.15, corrected). As expected, reconstruction was not above chance in M1 (*P* = 0.64, corrected). Notably, reconstruction in LOTC was not significant when images were not recognized at E2 (permutation test: *P* = 0.90, uncorrected). Thus, we observed robust LOTC-based reconstruction of E1’s temporal context, but only when subjects had the phenomenological experience of remembering a stimulus at E2. This reinstatement effect is striking when considering that subjects were only deciding whether each E2 stimulus was “old” or “new”; in other words, reinstatement of stimuli that neighbored E1 was entirely incidental to the task.

Behavioral studies have shown that when a list of words is recalled from memory, stimuli tend to be clustered according to their temporal proximity during encoding^28^. These temporal contiguity effects—which are a primary inspiration for theories of temporal context reinstatement—have a characteristic shape wherein recall clustering is strongest for stimuli that were immediately adjacent during encoding (± 1) and falls off dramatically for stimuli further apart during encoding (effects typically level-off for stimuli separated by ± 3 positions)^3,4,29^. To test for a similar function in our reinstatement effects, we ran additional reconstruction analyses that defined temporal context images as E1 ± 2 and, separately, as E1 ± 3. In contrast to the significant reconstruction effect in LOTC for E1 ± 1, reconstruction was numerically lower and not above chance at E1 ± 2 (*P* = 0.15) or E1 ± 3 (*P* = 0.57; permutation test). An analysis of variance (ANOVA) with lag to E1 as a factor confirmed a main effect of lag on reconstruction accuracy (**Fig. 2d**; *F*_2,14_ = 3.79, *P* = 0.048). Thus, reconstruction accuracy in LOTC decreased as a function of the temporal context images’ distance to E1, strongly mirroring temporal contiguity effects in free recall.

Finally, we tested whether temporal context reinstatement in LOTC was guided by the hippocampus. In prior work using this same fMRI dataset, we found that re-expression of hippocampal activity patterns across stimulus encounters predicted subsequent behavioral expressions of temporal memory (memory for when E1 occurred)^30^. While this finding is compatible with an account wherein the hippocampus reinstates past temporal contexts (thereby benefiting temporal memory), it is also open to other interpretations.

To test whether the hippocampus guided temporal context reinstatement, we first ran linear mixed-effects models using stimulus-specific E1-E2 pattern similarity (in medial temporal lobe regions) to predict temporal context reconstruction accuracy in LOTC. These models controlled for additional factors that might co-vary with temporal context reinstatement (E1 onset, E1-E2 lag; see Methods). Among the medial temporal lobe ROIs we considered, only hippocampal subfields CA1 and CA2/3/dentate gyrus (CA23DG) predicted temporal context reinstatement in LOTC (**Fig. 3a**; CA1: *ß* = 0.024, *P* = 0.007; CA23DG: *ß* = 0.027, *P* = 0.005; linear mixed-effects models, uncorrected). Specifically, when hippocampal activity patterns were more consistent across stimulus encounters (greater E1-E2 similarity), LOTC expressed stronger reinstatement of the information that *surrounded* the initial stimulus encounter. Notably, reversed versions of this analysis (in which E1-E2 pattern similarity in LOTC was used to predict temporal context reconstruction accuracy in the hippocampus) did not reveal significant relationships (*P*’s > 0.30).

**Fig 3.**
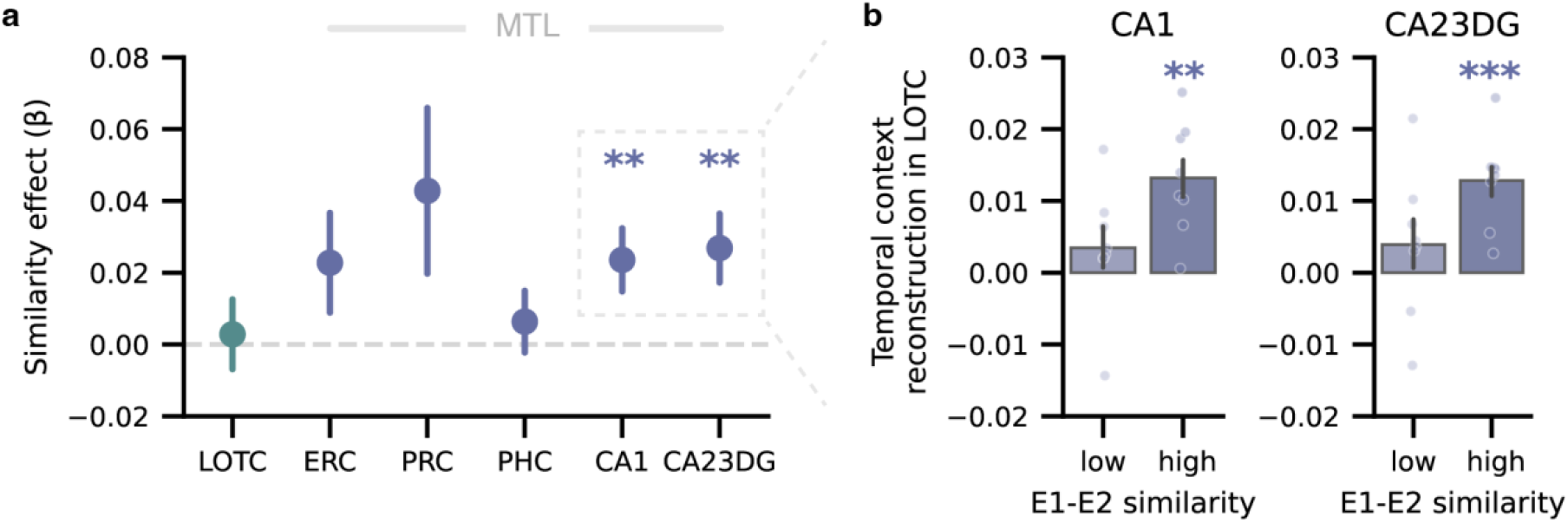
Hippocampus-mediated temporal context reconstruction. **a**, Temporal context reconstruction accuracy in LOTC was predicted by stimulus-specific pattern similarity (E1-E2 similarity) in hippocampal subfields CA1 and CA23DG (CA1: *ß* = 0.024, *P* = 0.007; CA23DG: *ß* = 0.027, *P* = 0.005; other ROIs: *P*s > 0.11; linear mixed-effects models, uncorrected). **b**, LOTC temporal context reconstruction accuracy was significantly greater than chance (zero) for images with high E1-E2 pattern similarity in the hippocampus (CA1: *t*_7_ = 4.75, *P* = 0.002; CA23DG: *t*_7_ = 5.56, *P* < 0.001; two-tailed one-sample *t* tests, uncorrected). Dots indicate individual subject data and bars represent the across-subject mean. In all panels, error bars depict s.e.m.; \*\**P* < 0.01; \*\*\**P* < 0.001.

As a complementary analysis, we binned images with high versus low E1-E2 hippocampal pattern similarity (subject-specific median splits). We found that LOTC reconstruction accuracy did not differ from zero for images with low CA1 or CA23DG similarity (**Fig. 3b**; *P*s > 0.30) but was significantly greater than zero for images with high similarity (CA1: *P* = 0.002; CA23DG: *P* < 0.001; two-tailed one-sample *t* tests, uncorrected). Reconstruction accuracy was also significantly greater for high vs. low similarity trials (CA1: *P* = 0.015; CA23DG: *P* = 0.025; permutation tests, uncorrected). Notably, E1-E2 similarity in LOTC did not predict temporal context reconstruction accuracy in LOTC (*P* = 0.77, uncorrected). Thus, the hippocampus and LOTC played complementary, but distinct roles in temporal context reinstatement. Specifically, our findings are consistent with the idea that the hippocampus guided cortical reinstatement (in LOTC)^31–33^.

## Discussion

In the present study, we used fMRI encoding models to reconstruct the temporal context in which events were originally encoded. Specifically, we show that when natural scene images are re-encountered, LOTC reinstates the visual content of *other scene images* that were encoded nearby in time to the original encounter. Importantly—and consistent with theoretical models—this temporal context reinstatement only occurred when events were successfully recognized and was strongest for scenes that were closest in time to the original event. Finally, the degree of temporal context reinstatement in LOTC was predicted by the strength of event-specific activity patterns in the hippocampus, revealing complementary roles for the hippocampus and high-level visual cortex in reinstating an event’s prior temporal context.

Our analysis approach and predictions were directly inspired by electrophysiological evidence demonstrating that remembering an event elicits reinstatement of neural activity patterns that were temporally-adjacent to that event^7–10,34^. However, our approach to measuring temporal context reinstatement is very distinct from these prior studies and affords more direct evidence of what, precisely, is reinstated. In particular, rather than testing whether neural activity patterns were reinstated, we tested whether the *content* of temporally-adjacent events (scenes) was reinstated. In fact, the encoding models we used were entirely ‘blind’ to the fMRI activity patterns elicited by E1 or temporally-adjacent events (E1 ± 1). Instead, the models only had access to the fMRI activity patterns elicited by E2. A subtle, but very important aspect of this approach is that it avoided the difficulty of teasing apart the neural activity patterns of E1 ± 1 from the neural activity pattern of E1. Teasing apart fMRI representations of temporally-adjacent events is fraught due to autocorrelation in the fMRI timeseries. In contrast, there is no autocorrelation in the *content* of temporally-adjacent scenes. Thus, by defining temporal context reinstatement in terms of scene content, we completely avoided the problem of fMRI autocorrelation and unambiguously demonstrate that the content of temporally-adjacent scenes was reinstated.

Another notable feature of our approach is that we did not test for reinstatement of a specific scene, per se, but instead tested for reinstatement of scene ‘features’^35–37^. That is, each scene was described in terms of a set of embeddings that corresponded to visual/semantic features. One of the appeals of this approach is that it has the potential to capture relatively subtle ways in which a given scene image (E1) incorporates information from temporally-adjacent scenes (E1 ± 1). For example, the representation of E1 may be imbued with specific content features such as animacy, affective qualities, physical locations, etc., from adjacent scenes^38^. Our approach is therefore amenable to identifying, at a granular level, the specific information that comprises a scene’s temporal context. A potential extension of this approach would be to test whether or how the specific features that comprise a scene’s temporal context influence the way in which the scene is remembered. For example, are some content features more likely than others to be incorporated into neighboring scenes?

Although we found that temporal context reinstatement was highly dependent on successful memory for E2, it is notable that subjects were only judging whether or not each scene was “old” or “new.” That is, subjects were not asked, or encouraged in any way, to encode associations between adjacent scenes or to try to remember these associations when making recognition memory decisions. Thus, temporal context reinstatement (operationalized here as reinstatement of temporally-adjacent scene content) was entirely incidental to the task demands. That said, even if subjects were not explicitly asked to remember temporal context information, one interpretation of the observed relationship between temporal context reinstatement and recognition memory success is that successful recognition may have occurred *because of* temporal context reinstatement^39,40^. In other words, temporal context reinstatement putatively provides memory with an episodic quality^41^.

Our findings also complement a substantial body of evidence implicating the hippocampus in encoding and remembering temporal information^17–19,42–51^. An emerging and important idea is that hippocampal activity patterns ‘drift’ over time^44^, with this drift potentially providing a direct window into changes in temporal context^52^. Here, we show that temporal context reinstatement in LOTC was predicted by the *similarity* of activity patterns in hippocampus across time (E1-E2 similarity). An appealing interpretation of this hippocampal similarity effect is that, even if hippocampal activity patterns drift over time, the hippocampus has the ability to ‘jump back in time’ to the original encounter^8,9^. In fact, in a recent study using this same dataset^30^, we found that E1-E2 similarity in the hippocampus predicted temporal memory for E1 (i.e., memory for *when* E1 occurred). Thus, even if the hippocampus is not directly reflecting reinstatement of scene *content*, our findings are consistent with the idea that the hippocampus tracks temporal information and plays a key role in guiding the process of temporal context reinstatement (expressed here in LOTC). The division of labor that we observe between the hippocampus and LOTC is also conceptually similar to other examples (outside the domain of temporal context reinstatement), where hippocampal and neocortical regions have been shown to play dissociable—but coupled—roles in memory reinstatement ^20,31– 33,53,54^.

Collectively, our findings provide unique evidence that remembering a stimulus elicits incidental reinstatement of the temporal context in which the stimulus previously occurred. In contrast to prior human studies which focused on reinstatement of neural activity patterns, we leveraged fMRI encoding models to explicitly show reinstatement of visual content from temporally-adjacent events. This finding strongly validates temporal context theories which argue that temporally-adjacent stimuli become part of a stimulus’ temporal context^1,55^. Moreover, by specifically linking trial-by-trial measures of cortical reinstatement to pattern similarity within the hippocampus, we also identify a division of labor wherein the hippocampus contains stimulus-specific codes that guide reinstatement of high-level visual information in neocortex.

## Methods

### Overview

All analyses were performed using the Natural Scenes Dataset (NSD), a large-scale dataset of high-resolution (7T) fMRI data from eight subjects who each performed a continuous recognition task on thousands of unique natural scene images over the course of 30–40 scan sessions. The results reported in this study are based on data from all of these sessions, from all subjects who participated in the NSD study. A detailed description of the dataset is reported in the original data publication^22^. Below we describe the specific methods relevant to the current study.

### Dataset

#### Subjects

Eight subjects (six females and two males; 19-32 years old) recruited from the University of Minnesota community participated in the NSD study. All subjects were right-handed with no known cognitive deficits or color blindness and with normal or corrected-to-normal vision. Subjects were not involved in the design or planning of the NSD experiment. Informed written consent was obtained from all subjects before the start of the study, and the experimental protocol was approved by the University of Minnesota Institutional Review Board.

#### Stimuli

All images used in NSD were drawn from the Microsoft Common Objects in Context (COCO) database^56^. A total of 73,000 unique images of natural scenes were prepared with the intention that each subject would be exposed to 10,000 distinct images (9,000 unique images and 1,000 shared images across subjects) three times each across the 40 scan sessions. Image presentations were pseudo-randomly distributed across sessions over the course of almost a year. The presentation structure was determined in advance and fixed across subjects so that difficulty of the recognition task was roughly similar across subjects. To provide a sense of the overall experimental design, the mean number of distinct images shown once, twice, or three times within a session was 437, 106, and 34, respectively.

#### Design and procedure

A detailed description of the task design is reported in the original data publication^22^. Briefly, subjects performed a continuous recognition task in which they decided, on each trial, whether the current image had been seen at any previous point in the experiment (“old”) or if they had not seen it before (“new”). Subjects completed up to 40 scan sessions in which they viewed up to 10,000 different color natural scene images. Each scene image was presented three times (if subjects completed all 40 sessions), with presentations pseudo-randomly spaced over the course of the entire experiment. Each scanning session consisted of 12 runs (750 trials per session). Each trial consisted of the presentation of an image for 3 s followed by a 1-s gap. Subjects were able to respond during the entire 4-s period and were also permitted to make multiple responses per trial (if they changed their mind). Because trials with multiple responses potentially captured a mixture of cognitive operations / decision making, we opted to exclude these trials from our analyses.

Four of the subjects completed the full set of 40 NSD scan sessions. Due to constraints on subject and scanner availability, two subjects completed 30 sessions, and two subjects completed 32 sessions. Accordingly, each subject viewed a total of 9,209-10,000 unique natural scene images across 22,500-30,000 trials.

#### fMRI data acquisition and preprocessing

MRI data was collected at the Center for Magnetic Resonance Research at the University of Minnesota. Imaging was performed on a 7T Siemens Magnetom passively-shielded scanner with a single-channel-transmit, 32-channel-receive RF head coil. Functional images were acquired using whole-brain gradient-echo echo-planar imaging (EPI) at 1.8-mm resolution and 1.6-s repetition time.

Details of the preprocessing of anatomical and functional data are reported in the original data publication^22^. Briefly, functional data were preprocessed by performing one temporal resampling to correct for slice time differences and one spatial resampling to correct for head motion within and across scan sessions, EPI distortion, and gradient non-linearities. Informed by the original data publication, the current study used the 1.0-mm volume preparation of the functional time-series data and “version 2” of the NSD single-trial betas.

### Data Analysis

The current study focused on the first two encounters with each stimulus (1st encounter = E1, 2nd encounter = E2). Our main analyses were restricted to E1/E2 stimuli that were (1) correctly identified as “new” at E1 (i.e., correct rejection) and correctly recognized as “old” at E2 (i.e., hit) and (2) were encountered in the same fMRI session, but in different scan runs. This resulted in a total of 721-1,875 stimuli that were used in the main analyses for each subject.

#### Regions of interest

Cortical ROIs, including bilateral angular gyrus (AG), lateral occipitotemporal cortex (LOTC), early visual cortex (V1), and primary motor cortex (M1) were drawn from the surface-based Human Connectome Project multimodal parcellation (HCP-MMP) atlas of human cortical areas^57^. All medial temporal lobe (MTL) subregions were manually drawn on the high-resolution T2 images obtained for each subject. These MTL ROIs included two subfields of the hippocampus (CA1 and CA2/3/dentate gyrus), along with entorhinal cortex (ERC), perirhinal cortex (PRC), and parahippocampal cortex (PHC).

#### Temporal context reconstruction analysis

To measure temporal context reinstatement (**Fig. 1**), we first extracted visual features of NSD scene images from the image encoder in OpenAI’s pretrained CLIP model with vision transformer backbone (ViT-32)^23^. We used the PyTorch implementation of this model. We then performed principal component analysis (PCA) on the extracted image embeddings to reduce the dimensionality of the visual features^35^. The first 20 principal components (PCs) were used to define the visual content of each image. For the primary temporal context reinstatement analyses, PC scores were averaged across E1-1 and E1+1 and these average scores were used in the encoding models.

To learn a direct linear mapping from the image to neural activity, we built voxel-wise encoding models to predict fMRI responses from image PC scores using cross-validated ridge regression (using scikit-learn Python package), separately for each subject and each ROI. The regularization parameter was determined by grid search over 11 values logarithmically spaced from 10–^5^ to 10^5^ to minimize prediction error. We then inverted this encoding model and applied it to held-out test data (leave-one-session-out) to reconstruct the image PC scores from fMRI activity patterns. Reconstruction accuracy was assessed by comparing the reconstructed and actual image PC scores using cosine similarity.

#### Pattern similarity analysis

Pattern similarity (within a given ROI) was calculated as the Pearson correlation between fMRI activity patterns evoked during different image encounters. Correlations were Fisher’s z-transformed before further analyses were performed. To avoid potential contamination from BOLD signal autocorrelation, all pattern similarity analyses were performed by correlating activity patterns for stimuli encountered across different runs (i.e., correlations were never performed within the same scanning run).

As in prior work^30^, we computed a stimulus-specific measure of E1-E2 pattern similarity. Specifically, for each target image, we computed “within-image” pattern similarity (E1 and E2 = same stimulus) and “across-image” pattern similarity (E1 and E2’ = different stimuli). The E2’ images used to compute across-image similarity were chosen from the same set of sessions (but different scanning runs) as the target image’s E1 and E2, thus controlling for differences in lag between image encounters. Further, E2’ images were chosen such that they had the same memory outcomes at the first two encounters as the target image. For target images with multiple possible E2’ images based on the criteria above, the median value of the across-image similarity scores was used. The across-image similarity was then subtracted from within-image similarity to yield a stimulus-specific measure of E1-E2 similarity for each image.

#### Statistics

Statistical analyses were performed using a combination of permutation tests, repeated-measures ANOVA, one-sample *t* tests, and mixed-effects regression models. A threshold of *P* < 0.05 was used to assess statistical significance.

To assess the statistical significance of reconstruction accuracy (**Fig. 2a, c**), we used non-parametric permutation tests (1,000 iterations). For each iteration of the permutation test, we shuffled the reconstructed image PC scores within each subject, then recomputed the average reconstruction accuracy (cosine similarity) across subjects. The 1,000 iterations yielded a null distribution of group-level reconstruction accuracy. We then computed one-tailed *P* values by determining how many samples from the null distribution exceeded the actual group-level reconstruction accuracy. For analyses shown in **Fig. 2a** and **2c**, *P* values were Bonferroni-corrected for multiple comparisons across 4 ROIs (AG, LOTC, V1, and M1).

To assess reconstruction accuracy as a function of temporal context images’ lag to E1 (**Fig. 2d**), permutation tests were performed within subjects (1,000 iterations) to obtain a subject-specific null distribution for each lag (E1 ± 1, E1 ± 2, E1 ± 3). The actual reconstruction accuracy for each subject, at each lag, was compared to the null distribution and expressed as a *z* score. Repeated-measures ANOVA was then used to test whether reconstruction accuracy (mean *z* score) varied by lag.

For analyses linking temporal context reconstruction accuracy to pattern similarity (**Fig. 3**), we used two approaches. The first approach (**Fig. 3a**) used mixed-effects regression models. Specifically, we ran mixed-effects linear regression models that predicted trial-by-trial reconstruction accuracy (cosine similarity) from stimulus-specific pattern similarity while including (controlling for) several additional factors. These additional factors of no interest included the lag between the beginning of the first trial in the experiment and E1 onset and the E1-E2 lag. All temporal lags were quantified in seconds and transformed with the natural logarithm to correct for non-normality in the distribution. All mixed-effects models included random intercepts per subject. The second approach (**Fig. 3b**) compared reconstruction accuracy for trials associated with high versus low pattern similarity. Specifically, we median split trials within each subject according to stimulus-specific pattern similarity (high vs. low) in a given ROI. We first compared reconstruction accuracy to chance (zero), separately for each bin (high vs. low pattern similarity), using two-tailed one-sample *t* tests.

Separately, we compared reconstruction accuracy between the two bins (high vs. low pattern similarity) using permutation tests (1,000 iterations). For each permutation, we randomly shuffled each subject’s pattern similarity labels across trials and computed the mean difference, across all subjects, in reconstruction accuracy for high vs. low pattern similarity trials. These permuted group-level differences served as the null distribution. Actual differences in reconstruction accuracy for high vs. low pattern similarity trials was expressed as one-tailed *P* values by determining how many samples from the null distribution exceeded the actual group-level difference. For these analyses, *P* values were not corrected for multiple comparisons because we had a strong *a priori* interest in the hippocampus (CA1 and CA23DG).

## Data availability

The data used in this study are openly available as part of the Natural Scenes Dataset, publicly available at http://naturalscenesdataset.org.

## Code availability

The code used for data analysis will be made available upon publication at https://github.com/futingzou/nsdTemporalContext.

## Acknowledgements

This work was supported by NIH-NINDS 2R01NS089729 to B.A.K.

